# The fecal microbiota of piglets is influenced by the farming environment and is associated with piglet robustness at weaning

**DOI:** 10.1101/859215

**Authors:** Diana Luise, Mathilde Le Sciellour, Arnaud Buchet, Rémi Resmond, Charlène Clement, Marie-Noelle Rossignol, Deborah Jardet, Olivier Zemb, Catherine Belloc, Elodie Merlot

## Abstract

This study describes the response of piglet microbiota to weaning in various environments, and investigates whether microbiota composition is associated with the robustness of piglets. Faecal samples and growth data were collected just before and 7 days after weaning in 288 piglets from 16 commercial farms characterised by their pathogenic status and antimicrobial use. The effect of weaning on the most abundant microbial families of faecal microbiota was roughly the same in all farms and in agreement with previous findings. Four enterotypes, ubiquitous in all farms, were identified, for which the most discriminating genera were *Prevotella*, *Faecalibacterium*, *Roseburia*, and *Lachnospira.* They corresponded to a gradual maturational shift from pre to post-weaning microbiota. Besides antimicrobial use, the pathogenic status of the farm had a major influence on the post-weaning microbial evolution of apparently healthy piglets. Regarding individual characteristics, piglets whose growth was less perturbed by weaning had more *Bacteroidetes* (P < 0.01) and less *Proteobacteria (P < 0.001).* In response to weaning, they showed a greater increase in *Prevotella* (P < 0.01), *Coprococcus* (P< 0.01) and *Lachnospira* (P < 0.05) than piglets that grew more slowly. Thus, the microbiota of robust piglets shares common characteristics regardless the living environment of animals.

## Introduction

In pig husbandry, weaning is a critical health-challenging period for piglets combining nutritional, environmental and social changes. Intestinal homeostasis plays a major role in maintenance of general health and prevention of infectious diseases (Bischoff, 2011). This is particularly relevant for piglets at weaning, for which the abrupt environmental changes may generate a dysbiosis, favourable to pathogenic bacteria causing post-weaning diarrhoea (PWD) (Gresse *et al.*, 2017). Even in the absence of PWD, weaning is associated with a transient reduction in growth due to a temporary impaired absorption of nutrients, often combined with a subclinical deterioration of animal health. Thus, in the absence of clinical signs, the robustness of piglets can be estimated by their post-weaning live weight trajectory (Revilla *et al.*, 2019).

The link between the microbiota, the feed intake and digestibility in one hand, and the health of the animal in the other hand, is still debated. In older pigs, gut microbiota appeared to be involved in feed efficiency (McCormack *et al.*, 2017; Le Sciellour *et al.*, 2018; Verschuren *et al.*, 2018), and it has been suggested that in weaner pigs also, the gut ecology could play a role in the differences in average daily gain (ADG) observed among piglets grown in an experimental farm (Mach *et al.*, 2015; Kiros *et al.*, 2019). Regarding health, individual differences in gut microbiome before weaning are linked to health disorders after weaning: animals experiencing PWD can be discriminated from healthy ones by their faecal microbiota as soon as one week after birth (Dou *et al.*, 2017). However, it is not known whether microbiota plays a role in piglet robustness in subclinical situations. One of the major issue is the dependence on the environment. Indeed, the impact of the rearing environment on the microbiota of piglets was shown for the genera *Oscillospira*, *Megasphaera*, *Parabacteroides*, and *Corynebacterium*, which differed among pigs from different farms (Yang *et al.*, 2018). In mice, it has been shown that observed links between microbiome composition and health are a highly variable across different facilities (Parker *et al.*, 2018). Furthermore, in commercial farms, the frequent and highly variable use of antimicrobials also interferes with these processes.

The aim of the present study was to explore the variability in piglet faecal microbiota around weaning across various environments and to link it with the robustness of the piglets. We collected faecal samples just before and 7 days after weaning in 288 piglets from 16 commercially-operating farms characterised by their pathogenic status and antimicrobial use (Table 1). Because individual daily health recordings are difficult to obtain in field studies based on high numbers of animals, the growth performance of the piglets was used as a proxy of their robustness.

**Table 1.**
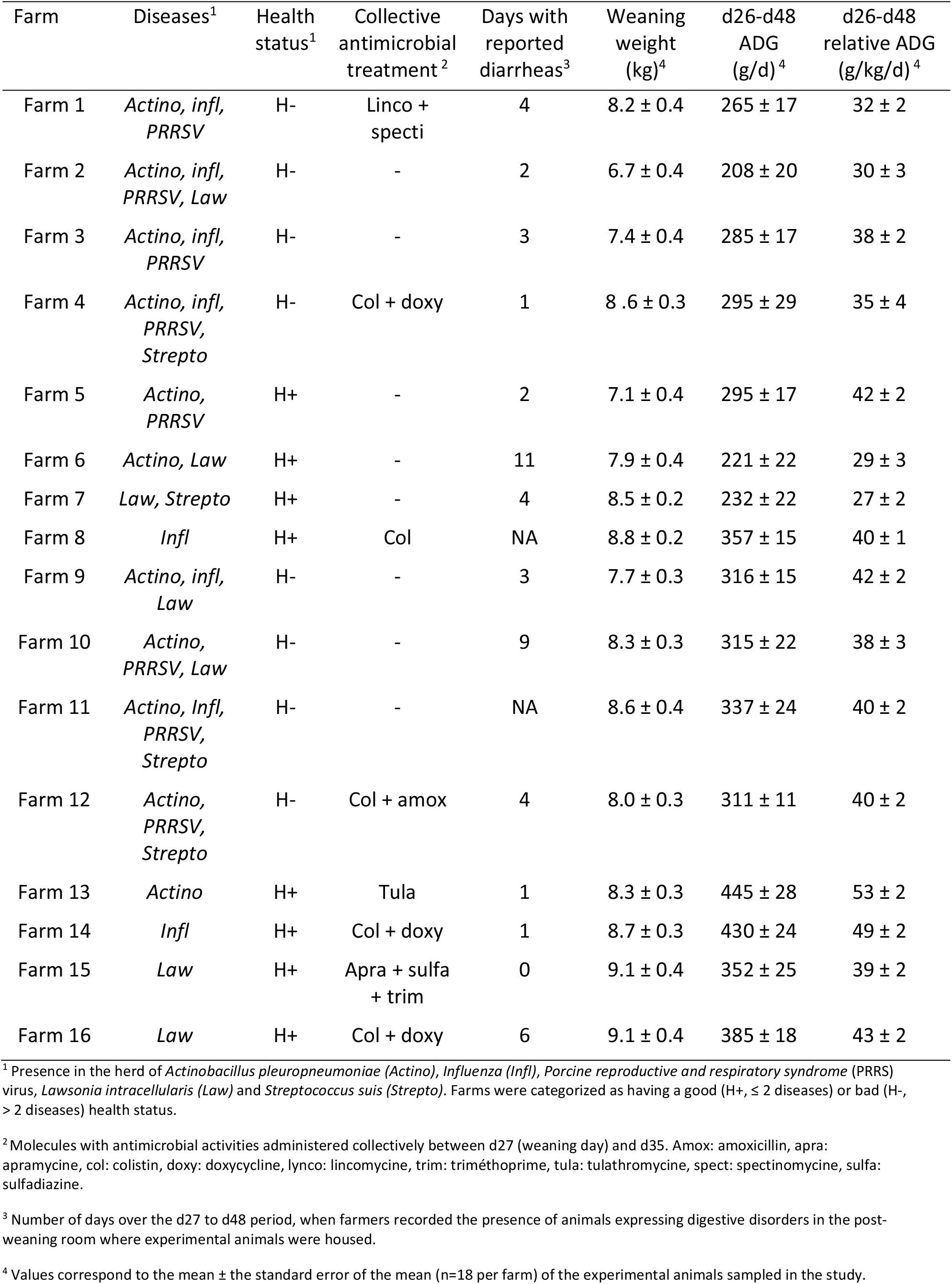
Characterisation of the health status of the 16 farms, and growth performances of the animals sampled around weaning in the study.

| Farm | Diseases<sup>1</sup> | Health status<sup>1</sup> | Collective antimicrobial treatment<sup>2</sup> | Days with reported diarrheas<sup>3</sup> | Weaning weight (kg)<sup>4</sup> | d26-d48 ADG (g/d)<sup>4</sup> | d26-d48 relative ADG (g/kg/d)<sup>4</sup> |
| --- | --- | --- | --- | --- | --- | --- | --- |
| Farm 1 | *Actino, infl, PRRSV* | H- | Linco + specti | 4 | 8.2 ± 0.4 | 265 ± 17 | 32 ± 2 |
| Farm 2 | *Actino, infl, PRRSV, Law* | H- | - | 2 | 6.7 ± 0.4 | 208 ± 20 | 30 ± 3 |
| Farm 3 | *Actino, infl, PRRSV* | H- | - | 3 | 7.4 ± 0.4 | 285 ± 17 | 38 ± 2 |
| Farm 4 | *Actino, infl, PRRSV, Strepto* | H- | Col + doxy | 1 | 8.6 ± 0.3 | 295 ± 29 | 35 ± 4 |
| Farm 5 | *Actino, PRRSV* | H+ | - | 2 | 7.1 ± 0.4 | 295 ± 17 | 42 ± 2 |
| Farm 6 | *Actino, Law* | H+ | - | 11 | 7.9 ± 0.4 | 221 ± 22 | 29 ± 3 |
| Farm 7 | *Law, Strepto* | H+ | - | 4 | 8.5 ± 0.2 | 232 ± 22 | 27 ± 2 |
| Farm 8 | *Infl* | H+ | Col | NA | 8.8 ± 0.2 | 357 ± 15 | 40 ± 1 |
| Farm 9 | *Actino, infl, Law* | H- | - | 3 | 7.7 ± 0.3 | 316 ± 15 | 42 ± 2 |
| Farm 10 | *Actino, PRRSV, Law* | H- | - | 9 | 8.3 ± 0.3 | 315 ± 22 | 38 ± 3 |
| Farm 11 | *Actino, Infl, PRRSV, Strepto* | H- | - | NA | 8.6 ± 0.4 | 337 ± 24 | 40 ± 2 |
| Farm 12 | *Actino, PRRSV, Strepto* | H- | Col + amox | 4 | 8.0 ± 0.3 | 311 ± 11 | 40 ± 2 |
| Farm 13 | *Actino* | H+ | Tula | 1 | 8.3 ± 0.3 | 445 ± 28 | 53 ± 2 |
| Farm 14 | *Infl* | H+ | Col + doxy | 1 | 8.7 ± 0.3 | 430 ± 24 | 49 ± 2 |
| Farm 15 | *Law* | H+ | Apra + sulfa + trim | 0 | 9.1 ± 0.4 | 352 ± 25 | 39 ± 2 |
| Farm 16 | *Law* | H+ | Col + doxy | 6 | 9.1 ± 0.4 | 385 ± 18 | 43 ± 2 |

<sup>1</sup> Presence in the herd of *Actinobacillus pleuropneumoniae* (*Actino*), *Influenza* (*Infl*), *Porcine reproductive and respiratory syndrome* (PRRS) virus, *Lawsonia intracellularis* (*Law*) and *Streptococcus suis* (*Strepto*). Farms were categorized as having a good (H+, ≤ 2 diseases) or bad (H-, > 2 diseases) health status.
<sup>2</sup> Molecules with antimicrobial activities administered collectively between d27 (weaning day) and d35. Amox: amoxicillin, apra: apramycin, col: colistin, doxy: doxycycline, lynco: lincomycin, trim: triméthoprim, tula: tulathromycin, spect: spectinomycin, sulfa: sulfadiazine.
<sup>3</sup> Number of days over the d27 to d48 period, when farmers recorded the presence of animals expressing digestive disorders in the post-weaning room where experimental animals were housed.
<sup>4</sup> Values correspond to the mean ± the standard error of the mean (n=18 per farm) of the experimental animals sampled in the study.

## Results

### Influence of weaning on piglet microbiota

After exclusion of low quality samples, the final dataset included data from 222 suckling (26-day old) and 254 weaned (35-day old) piglets, with a minimal number of 5003 reads per sample after rarefaction, and including 1 177 OTUs (Supplementary Table 1). *Firmicutes* and *Bacteroidetes* were the most abundant phyla at both ages and accounted for 63.0 % and 28.6 % of total sequences followed by *Proteobacteria* (5.2 %), *Spirochaetes* (1.7 %) and *Fusobacteria* (1.3 %). Both the observed and Shannon diversity indexes increased after weaning (P < 0.01, Table 2). Concerning the beta diversity, a variation of the composition of bacterial communities was observed according with the age of the animals and samples partially clustered for age (P < 0.001, Supplementary Fig. 1).

**Table 2.**
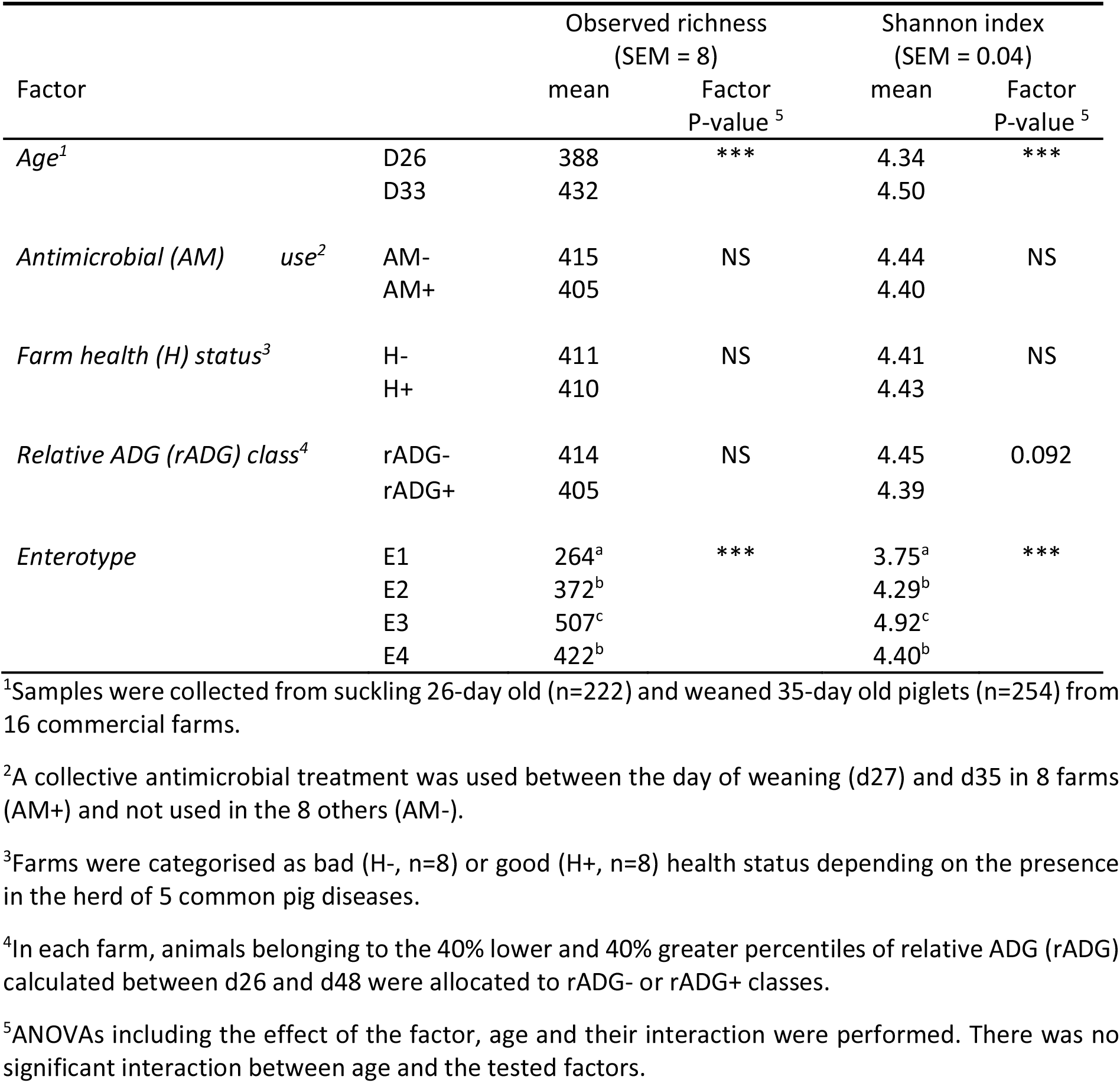
Influence on alpha diversity indexes of weaning, antimicrobial use, farm health status, growth aptitude of piglets and their enterotype belonging.

The comparison of the relative abundance of taxa indicated a clear effect of weaning, with 950 of the 1 177 OTUs having a fold-change significantly affected by age (FDR < 0.05, 514 decreased and 436 increased after weaning). Theses OTUs belonged to 34 families (Supplementary Fig. 2). Among these families, 16 had decreased and 8 had increased relative abundances after weaning (P < 0.05, Fig. 1). *Bacteroidaceae* (mainly *Bacteroides* genus) decreased by 61% after weaning (P < 0.001), including the species *B. fragilis, B. plebeius* and B. *uniformis*. *Christensenellaceae* (−35%*, P < 0.001)*, *Enterobacteriaceae* (− 42%) and *Clostridiaceae* (−32%, P < 0.001) decreased after weaning. *Prevotellaceae (*mainly *Prevotella)* increased by + 143% (P < 0.001). The abundance of *Lachnospiraceae* globally increased (+21%, P < 0.001), and especially *Blautia* (+69%), *Lachnospira* (+125%) and *Roseburia* genera (+150%, P < 0.001). The global abundance of *Ruminococcaceae* was stable along time, but in this family, *Faecalibacterium* increased after weaning (+117%, P < 0.001, and in particular *F. Prausnitzii*), *Oscillospira* decreased (−14%, P < 0.05), and *Ruminococcus* decreased (−22%, P < 0.05, especially *R. Gnavus)*. Some other less abundant families displayed important variations of abundance compared to d26 such as *Porphyromonodaceae* (−52%), *Fusobacteriaceae* (−62%), *Enteroccaceae* (−93%), *Rikenellaceae* (−81%), *Succinivibrionaceae* (+87%), *Veillonellaceae* (+173%) and *Peptostreptococcaceae* (+100%).

**Figure 1.**
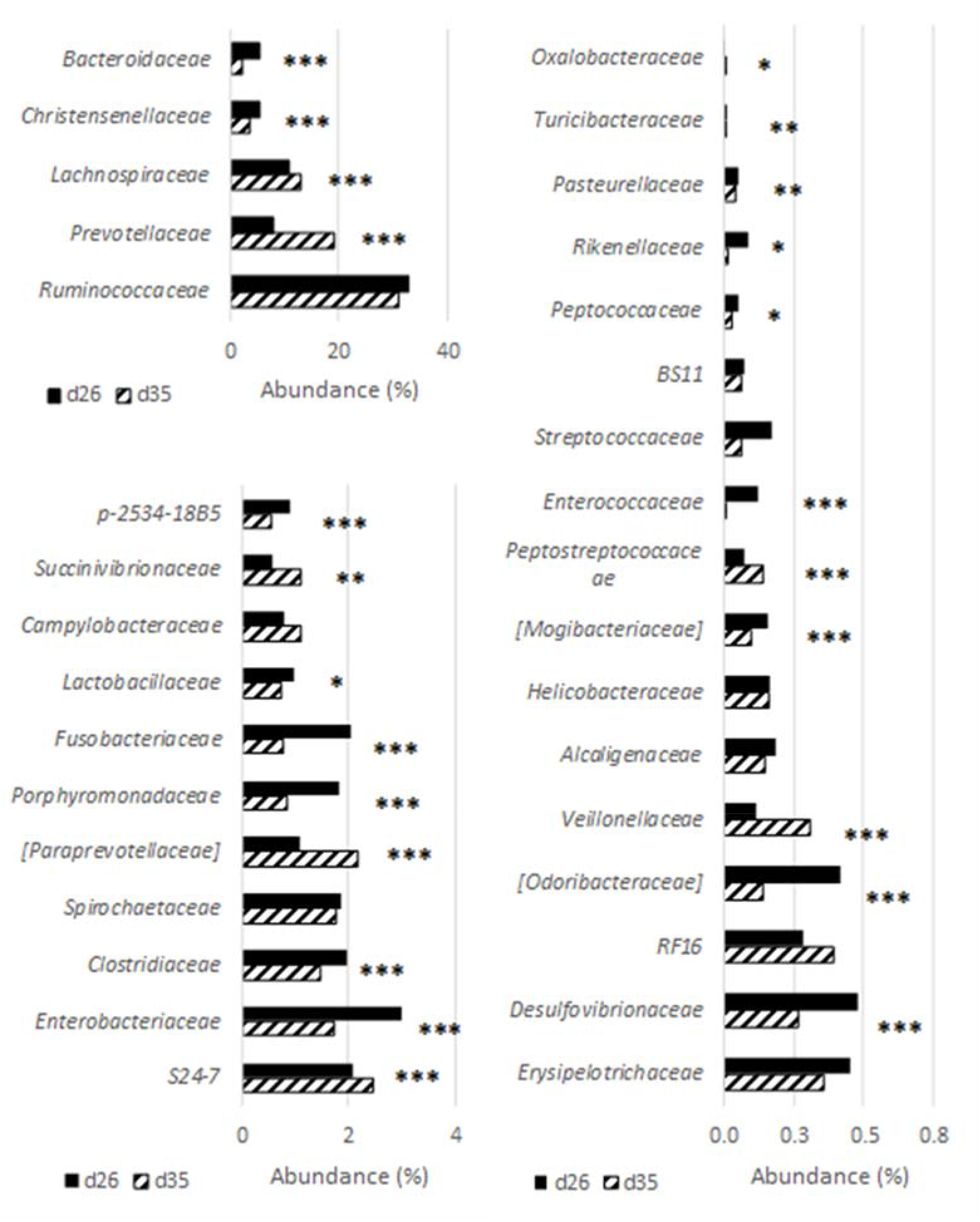
Relative abundance of the 33 families detected in piglet faecal samples before and after weaning. Samples were collected from suckling 26-day old (n=222) and weaned 35-day old piglets (n=254) from 16 commercial farms. For each taxon, a significant effect of time is indicated by *P <0.05, ** P <0.01, * P <0.05.

### Determination of enterotypes

Four enterotypes (E1 to E4) were established (Fig. 2A) using the methodology of Arumugam *et al.* (Arumugam *et al.*, 2011), for which the most discriminating genera were *Prevotella*, *Faecalibacterium*, *Roseburia*, and *Lachnospira* (Table 3). Both E1 and E2 showed a lower abundance in these four genera compared to E3 and E4. The E4 had the highest relative abundance in these four genera. Based on the number of OTU and the Shannon index, the diversity increased between the enterotypes from E1 to E3 (P < 0.001, Table 2). Between d26 and d35, 75% of the piglets shifted from an enterotype to another one (Fig. 2B). This dynamical shift seemed to evolve gradually from E1 to E2, to E3, and to E4.

**Figure 2.**
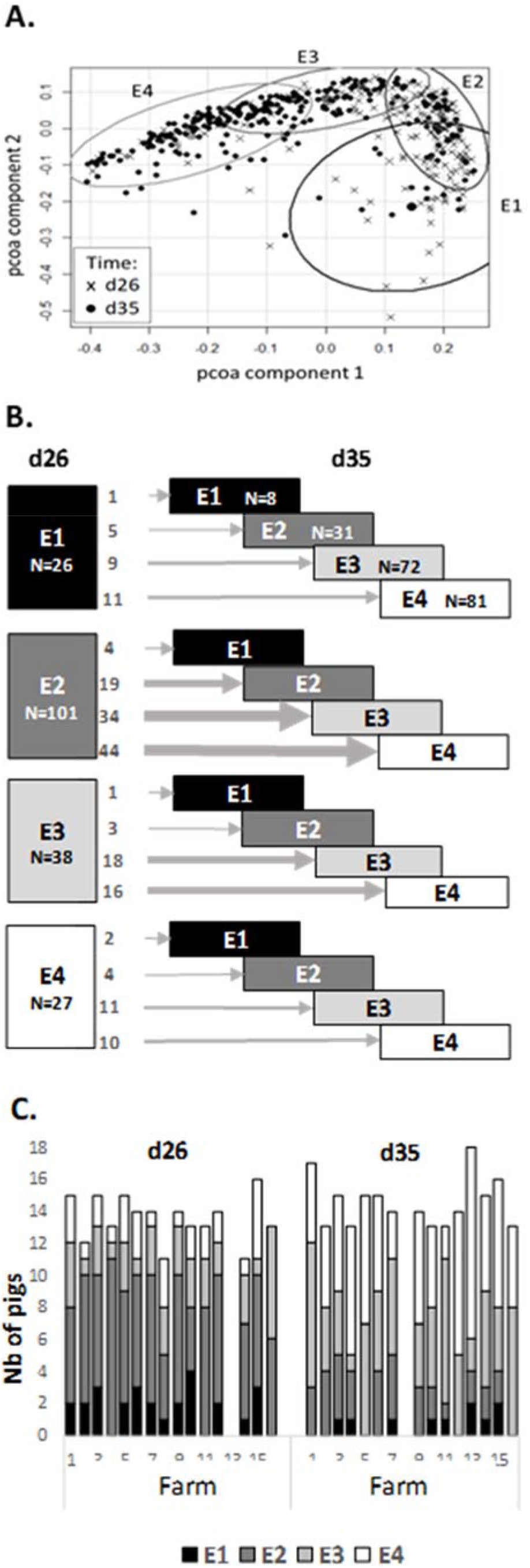
Identification of four enterotypes in faecal samples and their distribution in pre- and post-weaning piglets. Four enterotypes named E1 to E4 could be identified among the samples collected from suckling 26-day old (n=222) and weaned 33-day old piglets (n=254) in 16 commercial farms (A). The 192 pigs with data available at both time points were used to show the number of pigs in each enterotype at d26 and their new enterotype allocation at d35 (B). The variability of their distribution between farms is illustrated by the distribution of enterotypes at d26 and d35 (C).

**Table 3.** Relative abundance of genera discriminating the E1, E2, E3 and E4 enterotypes.

| Genus<sup>1</sup> | Mean relative abundance (%) |  |  |  | Enterotype effect (P-value)<sup>2</sup> |
| --- | --- | --- | --- | --- | --- |
|  | E1 | E2 | E3 | E4 |  |
| *Prevotella* | 3.87a | 1.32a | 11.29b | 31.4c | <0.001 |
| *Faecalibacterium* | 0.35a | 0.24a | 2.41b | 11.08c | <0.001 |
| *Roseburia* | 0.24a | 0.17a | 1.20b | 4.36c | <0.001 |
| *Lachnospira* | 0.02a | 0.05a | 0.48b | 1.08c | <0.001 |
| *Bacteroides* | 16.10a | 5.71b | 0.59c | 0.63d | <0.001 |
| *Coprococcus* | 0.12a | 0.35b | 0.86c | 0.93c | <0.001 |
| *Treponema* | 0.54a | 1.21b | 3.77c | 1.09b | <0.001 |
| *Paludibacter* | 0.02a | 0.26b | 0.27c | 0.06a | <0.001 |
| *Ruminococcus* | 1.27a | 2.25b | 2.16c | 1.34a | <0.001 |
| *Mitsuokella* | 0.02ab | 0.01a | 0.03b | 0.21c | <0.001 |
| *Oscillospira* | 2.43a | 3.59b | 2.79c | 2.05a | <0.001 |
| *Lactobacillus* | 1.82a | 0.88bc | 0.78c | 0.68d | <0.001 |
| *Campylobacter* | 1.52a | 0.57b | 1.14a | 1.11a | <0.001 |

<sup>1</sup>The table presents a non-exhaustive list of genera that were differently expressed among enterotypes. Means were calculated using the total dataset, from suckling 26-day old (n=222) and weaned 35-day old piglets (n=254) from 16 commercial farms.
<sup>2</sup>The order of genera in the table follows the P-value of the Kruskal-Wallis test for the enterotype effect (greatest for *Prevotella*, lowest for *Campylobacter*).

### Influence of farm environment on piglet microbiota

At both d26 and d35, the richness varied among farms but not the Shannon index (P < 0.05 and > 0.1, respectively). The effect of weaning on most abundant microbial families was roughly the same in all farms (Supplementary Fig. 3). However, there was a significant variability associated to the farm. Before weaning (d26), farms differed for the abundance of *Christensenellaceae*, which was the 5^th^ more abundant family in piglet microbiota, and for 5 other less abundant families, including *Lactobacillaceae* (P< 0.05, Supplementary Table 2). After weaning (d35), the differences due to the environment increased, and farms differed in their relative abundance for 21 families (P < 0.05). The 4 enterotypes were observed in all farms, at least at one of the two different ages (Fig. 2C), but nevertheless the repartition of enterotypes was heterogeneous among farms at d35 (P < 0.05).

A health score ranging from 0 to 5 was attributed to farms according to the presence or absence of five major swine pathogens (*Actinobacillus pleuropneumoniae*, *Influenza*, Porcine reproductive and respiratory syndrome virus, *Lawsonia intracellularis* and *Streptococcus suis)*. Farms hosting 1 or 2 of these diseases were considered to display a good sanitary status (H^+^, 8 farms); farms hosting 3 or more of diseases were assigned to the bad sanitary status group (H−). The dataset included information for 101 and 120 H+ samples at d26 and d35 respectively, and for 121 and 134 H− samples at d26 and d35 respectively. Farm sanitary status did not influence the alpha diversity indices (Table 2) nor the distribution of piglets among E1 to E4 enterotypes (data not show). A variation of the composition of bacterial communities (beta diversity) was observed according with the sanitary status of the farm (P = 0.04). The H+ animals differed from H− for 436 OTUs at d26 (P < 0.05, 103 increased and 333 decreased in H−), for 340 OTUs at d35 (P < 0.05, 203 increased and 137 decreased in H−, Supplementary Fig. 4), and 152 of these OTUs were common to both ages. These OTUS belonged to 27 different families. Independently of age, H− piglets tended to have more *Paludibacter* (P < 0.1) than H+ piglets (Fig. 3A). The weaning-associated decrease of *Oscillospira* was more pronounced in H− pigs than H+ pigs (−22% vs. −5%, P < 0.05) and H− pigs had a higher abundance of *Helicobacteraceae* family and *Succinivibrio* genus at d35 than H+ pigs (+134%, P < 0.05 and +111%, P < 0.05, respectively). At d35, compared to H+, H− tended to have less *Ruminococcaceae* and more *Christensenellaceae* (−2.51% and +39%, P < 0.1). At species level, at d35, *Bacteroides uniformis* as well as *Enterococcus cecorum* were more abundant in H− than H+ pigs (mean LogFC of 0.96 and 0.66 respectively, P < 0.001).

**Figure 3.**
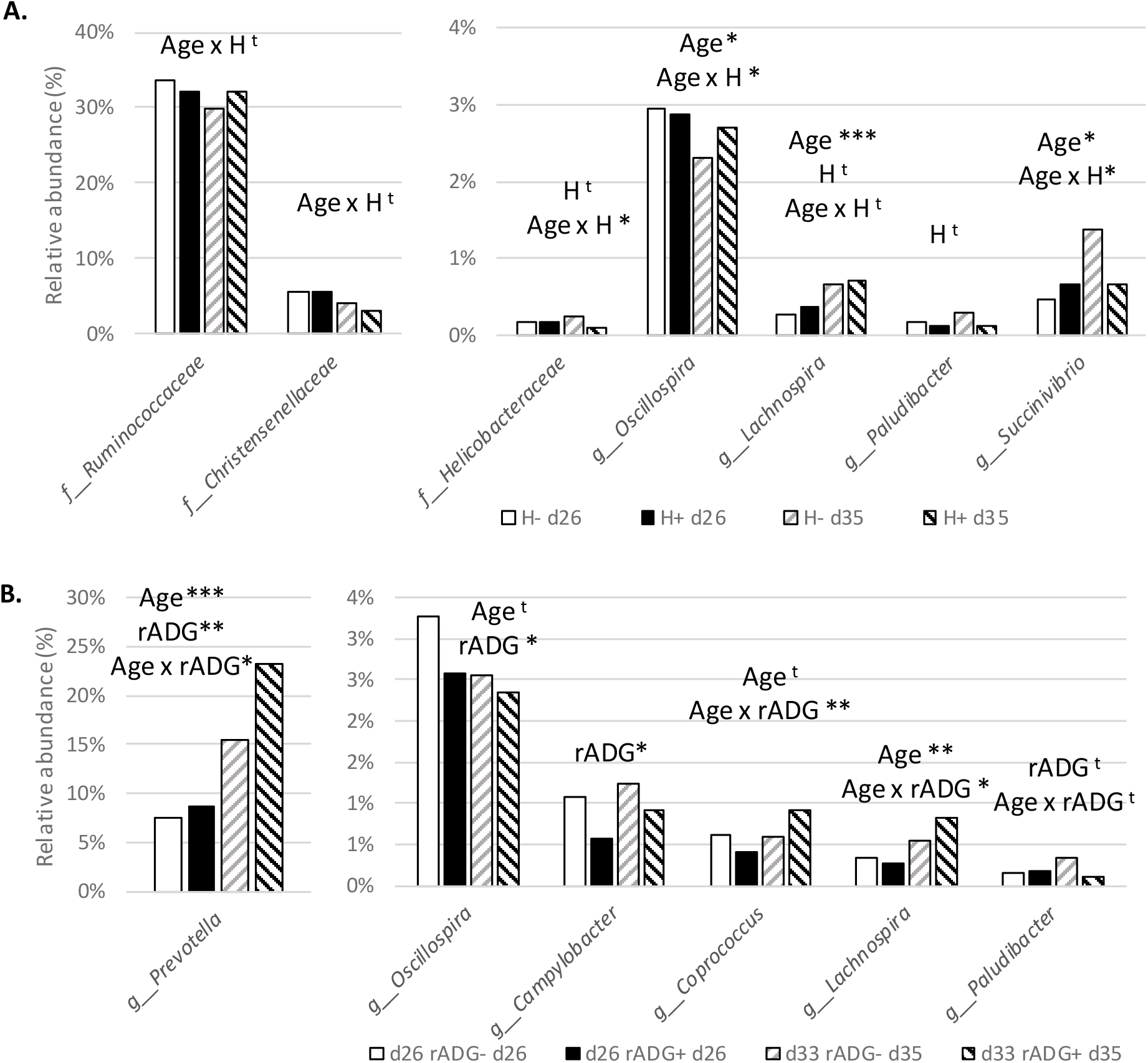
Bacterial taxa discriminating piglets from farms of contrasted health status and with different growth speed. Concerning the farm health status (A), the relative abundances of genera and families of bacteria differing among piglets raised in farms of low (H−) and good (H+) health status were determined from fecal samples collected at 26 (n=98 and 118 respectively) and 35 days of age (n=119 and 130, respectively). The relative abundances of taxa differing among animals showing a high (rADG+) and a low (rADG−) relative average daily gain compared to their farm-mates are presented (B). For each taxon, tendencies and significant effects of Age, Health status of the farm (H) or rADG class and the Age x H or rADG x Age interactions are indicated. *P <0.05, ** P <0.01, * P <0.05, t P<0.1.

An antimicrobial was administered during the first week after weaning in eight farms (AM+) while no antimicrobials were used during that period in the eight other farms (AM−, Table 1). Antimicrobial use did not influence the alpha diversity indices (Table 2) nor the distribution of piglets among E1 to E4 enterotypes, although a variation of composition of bacterial communities (beta diversity) was observed (P <0.01). The abundance of 248 OTUs was decreased (LogFC < 0, P < 0.05) and of 214 OTUs increased (LogFC > 0, P < 0.05,) in AM+ compared to AM− animals (Supplementary Fig. 5).

### Relation between piglet growth after weaning and microbiota composition

Growth performance of the pigs from the 16 farms is shown in Table 1. Piglets are fast growing animals for which the ADG over a given period of time is highly correlated to the body weight at the beginning of the considered period. To analyse piglet growth independently of its pre-weaning life trajectory, and thus to consider only its robustness to weaning, the relative ADG (the ratio between ADG over the d26 to d47 period and the weight at d26) was calculated. As expected, weight at weaning, ADG and rADG were highly variable among farms (P < 0.001). AM+ pigs had greater ADG and rADG (P < 0.001, data not shown). Pigs raised in H+ farms had a greater weight at weaning (8.5 vs 8.0 ± 0.2 kg, P < 0.05), a greater ADG (362 vs. 332 ± 11 g/day, P <0.001) but a comparable rADG to H− pigs (P > 0.1). Regarding enterotypes, compared to piglets that had reached the E4 enterotype 7 days after weaning, piglets at the E2 enterotype and E3 enterotypes had or tended to have a lower rADG (E4: 43.5 ± 1 vs. E2: 37 ± 2 g/kg/day, P < 0.05 and vs. E3: 40 ± 1 g/kg/day, P < 0.1). E1 pigs had an intermediate rADG (42 ± 3 g/kg/day).

In each farm, piglets showing the greatest rADG (top 40% of the farm distribution of the 18 sampled piglets) and the lowest rADG (bottom 40% of the farm distribution) were assigned to the rADG+ and rADG− classes (151 pigs with data for both time points). The rADG− class tended to have a higher Shannon index than the rADG+ class (P< 0.1) while the observed richness was similar in both classes (Table 2).

Pigs in the rADG− group differed from pigs in the rADG+ group for 315 OTUs at d26 (P < 0.05, 150 were decreased and 165 increased in rADG−), and for 461 OTUs at d35 (P < 0.05, 244 were decreased and 217 increased in rADG−), distributed in 14 and 15 different families respectively (Supplementary Fig. 6). At phylum level, whatever the age, rADG+ pigs had more *Bacteroidetes* (31.5 vs. 27.2%, P< 0.01) and less *Proteobacteria* (4.18 vs. 6.65%, P<0.001) than rADG− pigs. Differences were also detectable at the genus level: rADG+ pigs had less *Oscillospira* (P < 0.05) and *Campylobacter* (P < 0.01) than rADG− pigs (Fig 3B). There was a time x rADG group interaction for *Prevotella* (P < 0.01), *Coprococcus* (P< 0.01) and *Lachnospira* (P < 0.05), whose abundance increased after weaning, and for which the increase was even more visible in rADG+ pigs. The abundance of *Paludibacter* tended to be influenced by the time x rADG group interaction (P < 0.1).

The absolute value of rADG between d26 and d48 could be predicted by a linear model using data from microbiota at d26 and d35 with an adjusted R^2^ of 0.29 (Table 4). The model included: the abundance of the *Porphyromonodaceae* family at d26 and d35 (P < 0.001 and < 0.05), abundance of three genera (*Campylobacter* at d26, P < 0.01; *Fusobacterium*, P < 0.001; *Clostridium* at d35, P < 0.05), and of 4 OTUs belonging to *Ruminococcaceae* (OTU 4315785 at d26, P < 0.05), *Coprococcus* (OTUs 295683 and 752354 at d35, P < 0.001 and 0.05), and *Prevotella* (OTU 350627 *Prevotella copri* at d35, P < 0.001). As a comparison, the prediction of d26-d48 rADG by weight at d26 alone presented R^2^ of 0.03.

**Table 4.**
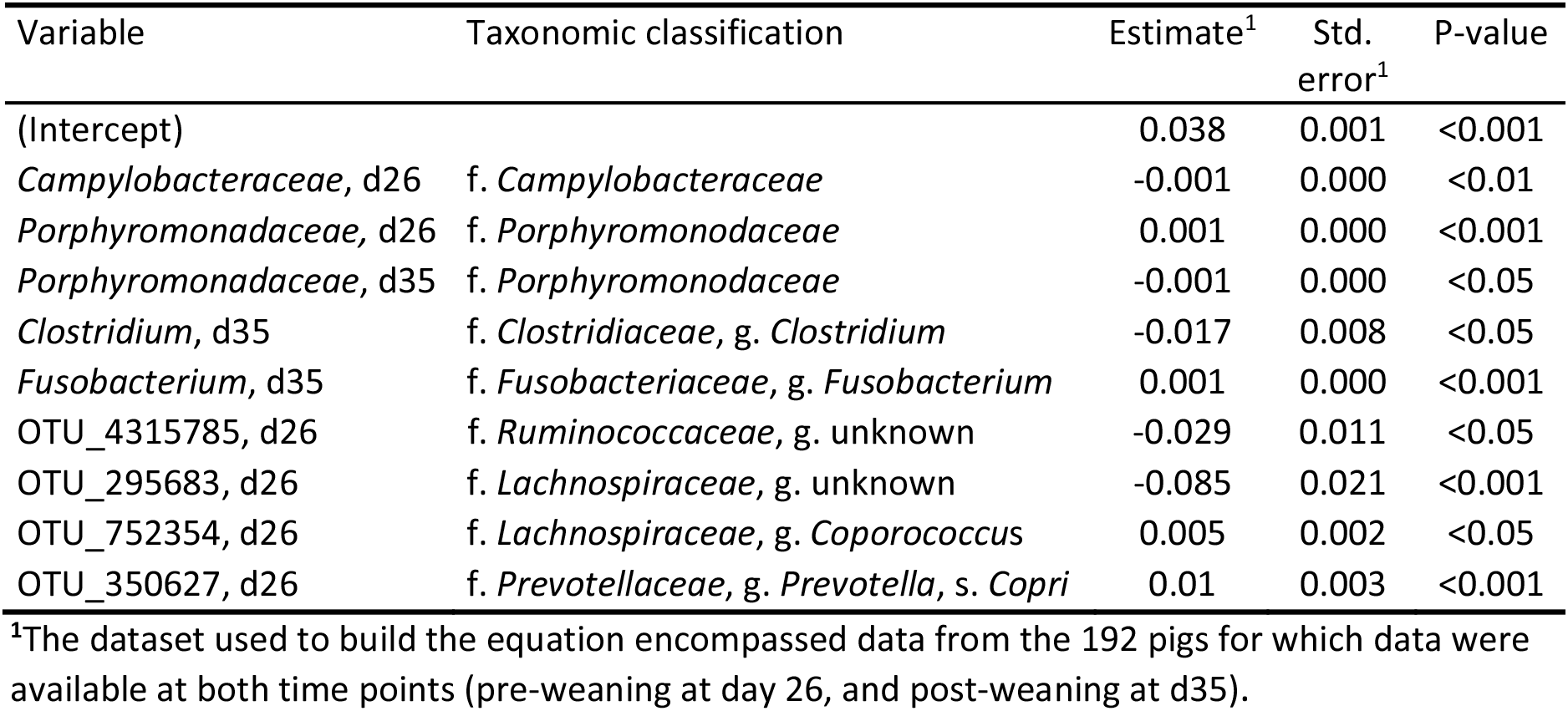
Estimates of the parameters of the linear model calculating rADG over the d26 - d48 period (R^2^ = 0.29).

## Discussion

### Microbial maturation and enterotypes around weaning

Our study had the particularity to investigate the variability of the effect of weaning on piglet microbiome in 16 different commercial farms simultaneously and on 222 pre- and 254 post-weaning piglets respectively. This allowed testing the universality of the changes previously reported in the literature (Mach *et al.*, 2015; Chen *et al.*, 2017; Motta *et al.*, 2019). Results from the present study show that weaning transition induced evident changes in the bacterial community. The alpha diversity values increased in the post-weaning comparing with the pre-weaning period, which is recognised as an indicator of greater stability and microbial maturity (Holman and Chenier, 2014; Chen *et al.*, 2017). A decrease of the abundance of *Enterobacteriaceae, Bacteroidaceae, Clostridiaceae, Enterococcaceae, Fusobacteriaceae, Porphyromonodaceae and Rikenellaceae* families and *Bacteroides*, *Fusobacterium, Ruminococcus* genera as well as an increase in *Prevotella*, *Veillonellaceae* and *Succinovibrionaceae* was observed after weaning. It is notable that this recognised microbial shift (Guevarra *et al.*, 2019) was observed despite the fact that pre and post weaning time points were close (d26 and d35), indicating a quick shift of the bacterial community.

It has been proposed that enterotype-like clusters could be described in pig microbiome, similarly to what has been reported in human populations (Arumugam *et al.*, 2011). Mach *et al.* (2015) identified that maturing bacterial communities in piglets evolved after weaning toward two different clusters, primarily distinguished by their abundances in unclassified *Ruminococcaceae* and *Prevotella*. Pre-weaning pigs all pertained to the *Ruminococcaceae* cluster and the shift toward the *Prevotella* cluster started by weaning, but the two phenotypes persisted in older fattening pigs (Ramayo-Caldas *et al.*, 2016). We identified four enterotype-like clusters around weaning, that were ubiquitous since they could be observed across the 16 farms. The shift from E1 to E4 was associated with an increase in the abundance of *Faecalibaterium*, *Lachnospira*, *Prevotella* and *Roseburia genera*, which were more abundant after than before weaning. Thus, E1 to E4 enterotypes probably reflect steps of increasing maturity of piglet microbiota from the suckling phase to a more mature microbiota able to use the new nutritive substrates provided by a cereal-based diet. According to their abundances of *Prevotella*, *Ruminococcus*, *Mitsuokella* and *Treponema*, the four enterotype-like clusters observed in the present study could be four sub-classes of the “*Ruminococcus* and *Treponem*a” enterotype described previously (Ramayo-Caldas *et al.*, 2016). This is coherent with the age of the pigs and the very short time elapsed since weaning, since the shift toward the “*Prevotella*” phenotype develops slowly after weaning (Mach *et al.*, 2015).

### Inter-farm variability of the microbial profile of piglets

In addition to the animal age, structure and activity of gut microbiota can differ between animals depending on the breed and genotype (Kubasova *et al.*, 2018; Luise *et al.*, 2019), diet (Le Sciellour *et al.*, 2018), health status (Bearson *et al.*, 2013) and antimicrobials (Holman and Chenier, 2014; Looft *et al.*, 2014; Soler *et al.*, 2018). The housing environment plays also an important role in shaping the gut microbiota. This has been demonstrated by comparing the gut microbiota of piglets raised indoor or outdoor (Mulder *et al.*, 2009), mice raised in a clean environment or in the presence of dust and soil (Zhou *et al.*, 2016), and human babies exposed to different house dusts (Konya *et al.*, 2014). In the present study, 16 commercial farms were included. Despite relatively uniform diets, genotypes and herd size, the observed farm effect on microbial profile was expected because these farms differed for factors that may influence the gut bacterial community such as their pathogens status, antimicrobial use, temperature, air condition in the room, or animal density. Among suckling piglets, farms differed for the abundances of *Christensenellaceae*, and *Lactobacillu*s, which are common genera in the pig hindgut and are considered beneficial bacteria able to improve gut health or feed efficiency (McCormack *et al.*, 2017; Valeriano *et al.*, 2017). After weaning, farms differed one from another for the abundance of two thirds of their present families. In order to disclose the farm effect on faecal microbial profiles, farms were classified on their antimicrobial use and sanitary status. The antimicrobial treatment after weaning influenced the abundance of 39% of the OTUs of the dataset compared to untreated piglets, showing that it plays a major role in the observed inter-farm variability. However, antimicrobial effects are highly dependent on the molecule, and the diversity of molecules used in the farms of the present study makes difficult the interpretation of the results (Bosi *et al.*, 2011; Looft *et al.*, 2014).

### Influence of farm health status on the microbial profile of piglets

The influence of the farm health status, determined according to the presence or absence of five major swine pathogens (*Actinobacillus pleuropneumoniae*, *Influenza*, Porcine reproductive and respiratory syndrome virus, *Lawsonia intracellularis* and *Streptococcus suis)* was also investigated. The farm health status did not influence the alpha diversity indices, while the overall bacterial community structure was slightly affected, indicating that the proportion of specific taxa in the swine faecal microbiota rather than community membership was influenced. The number of OTU differently expressed between H− and H+ farms was greater before weaning (37%) than after (29%), with 13% of OTUs affected at both ages. These OTUs belonged to many different genera, with few consequences on the global abundance of these genera. Noticeably, H+ and H− pigs differed at d35 in their abundance of *Oscillospira*, a genus whose abundance was already reported to vary depending on farm origin (Yang *et al.*, 2018). H− pigs also presented more *Succinivibrio* and *Helicobacter*. Particularly, *Helicobacter* genus encompasses *Helicobacter suis*, that can induce growth retardation, as well as gastritis, gastric lesions and infections in pigs, humans and also other species (Park *et al.*, 2000; De Bruyne *et al.*, 2012; Péré-Védrenne *et al.*, 2017). Using a model of deterioration of environmental hygiene, another study showed that housing pigs in dirty rooms had limited effects on gut microbiota, but also increased the abundance of *Helicobater* (Kubasova *et al.*, 2018).

The mechanism by which the global health status of the herd influenced the gut microbiota of piglets is unclear. None of the piglets displayed any clinical expression of the five recorded diseases, which usually develop in older pigs. However, the PCR and serological tests performed on the piglets included in this experiment indicated that, in the farms diagnosed as positive for one of these pathogens, some of the piglets were already carriers of this pathogen three weeks after weaning (data not shown). This early carriage is in agreement with the literature (Robertson *et al.*, 1991; Papatsiros *et al.*, 2006; Tobias *et al.*, 2014; Ryt-Hansen *et al.*, 2019). The influence of the farm health status on piglet microbiota cannot be due to a competition of theses specific pathogens with commensal bacteria since none of them develops in the gut lumen. *Lawsonia* intracellularis has a tropism for intestinal cells, but it was also present in both H+ and H− farms (4 and 3 farms respectively).

Some indices suggest that the immune system of piglets from H− farms might have been more solicited since birth. First, the fact that some animals had already seroconverted three weeks after weaning indicates that their immune system has been activated by these pathogens early in their life. Second, we described elsewhere that H− animals had greater numbers of blood circulating leukocytes before weaning, including lymphocytes, as well a higher concentrations of serum total IgM, which both suggest a higher stimulation of their immune system (Buchet *et al.*, 2018). Third, H− piglets were lighter than H+ piglets at weaning, suggesting that their global health might have been worse than H+ pigs from birth. The greater solicitation of the immune system of the piglets precisely during the post-natal phase might have influenced the maturation of their immune system and diversification of their immune repertoire (Levast *et al.*, 2014). More, the pathogens circulating in the herd are also expected to shape the immunity of the mothers, and the repertoire of antimicrobial and natural IgA secreted in their milk. Differences in maternal milk IgA and piglet immune maturation between H− and H+ conditions could have influenced the IgA repertoire of the suckling piglets, which protect the gut lumen against pathogens but also regulate commensal microbiota (Salmon *et al.*, 2009).

### Association between growth and microbiota

The absolute growth (ADG) after weaning is related to animal robustness but is also highly constrained by the starting weaning weight. The use of relative ADG allows considering the post-weaning growth performance corrected from the pre-weaning growth trajectory, and thus can be used as a proxy of robustness to the weaning event. Interestingly, we found that 29% of the variability of rADG could be predicted by the relative abundance of a few families, genera and OTUs present in the microbiota around weaning, confirming a link between robustness at weaning and gut flora composition.

Classes of animals showing low and high rADG relatively to their pen-mates were used to look for characteristics of the microbiota that would be shared independently of the rearing environment among robust piglets. We observed that the animals having the highest relative ADG in their farm had higher abundance of *Bacteroidetes* and lower abundance of *Proteobacteria* than their pen-mates who grew more slowly. Increased abundance of *Proteobacteria* is considered as a potential indicator of gut dysbiosis (Shin *et al.*, 2015; Adhikari *et al.*, 2019), thus the most robust piglets might have a microbiota that would be more resistant to dysbiosis. At genus level, they had higher abundance of *Prevotella*, after weaning, which has been reported to be positively correlated with ADG (Mach *et al.*, 2015; Kiros *et al.*, 2019). This greater abundance in better growing animals could be related to the capacity of *Prevotella* to produce enzymes that degrade complex dietary polysaccharides, improving fiber digestibility and host feed efficiency (McCormack *et al.*, 2017; Le Sciellour *et al.*, 2018). The abundance of *Prevotella* has also been associated with resistance to PWD (Dou *et al.*, 2017). Piglets of the rAGDG+ class also had more *Coprococcus* and *Lachnospira*, and less *Oscillospira*, *Campylobacter* and possibly *Paludibacter* compared to their farm-mates. The relationship between these genera and robustness of animals has not been reported before. Interestingly, this phenotype shared characteristics with the E4 enterotype (more *Prevotella*, *Coporcoccus*, and *Lachnospira*). In agreement, animals displaying the E4 enterotypes 7 days after weaning had indeed a higher relative growth than E2 and E3. This suggests that a faster maturation of microbiota after weaning would be associated with a better robustness. However, our results only display associations and cannot prove any causal relationship.

## Conclusion

In this study, the previously described effect of weaning on faecal microbiota was confirmed in a variety of commercial farms and we observed the presence of ubiquitous enterotypes among farms reflecting maturational stages of microbiota from a young suckling to an older cereal-eating. This maturation might be associated to the robustness of piglets, since piglets whose growth is less perturbed by weaning shared common characteristics of their microbiota regardless the farm where they are raised, and reached just after weaning a phenotype close to the enterotype 4. This study also reveal a significant influence of the rearing environment on the post-weaning evolution of piglet microbiota, which, besides antimicrobial use, also relied on the general health status of the herd.

## Experimental procedures

### Experimental design and farm characterization

A total of 288 Pietrain × (Large White × Landrace) piglets were included in the study. Piglets were sampled from 16 commercial farms having more than 100 sows and belonging to the COOPERL cooperative, from January to June 2015. Farms used diverse but conventional starter and weaner diets, and weaned piglets at four weeks of age. In each farm, two days before weaning, 9 sows of different parities were chosen to reflect the demographic pyramid of the herd, and two apparently healthy and middleweight entire males per litter were included in the study (18 piglets per farm). Piglets were weaned at 27 ± 2 days of age. Farms were visited 3 times: just before weaning, one week and 3 weeks after weaning, when piglets were in average 26 (d26), 35 (d35), and 48 ± 2 day-old (d48).

The health status of the farms was evaluated according to the veterinary follow-up of the farm regarding the five main pathogens currently observed in French pig farms: *Actinobacillus pleuropneumoniae*, *Influenza*, *Porcine reproductive and respiratory syndrome* (PRRS) virus, *Lawsonia intracellularis* and *Streptococcus suis*. Farms of diverse health statuses were chosen: farms serologically positive and / or with symptoms for 1 or 2 of the diseases mentioned above were considered to display a good health status (H^+^, 8 farms); farms hosting 3 or more of these diseases were assigned to the bad health status group (H^−^, 8 farms, Table 1). The presence of these pathogens among piglets during the experiment was confirmed by specific PCR and serologic tests performed on all piglets three weeks after weaning (data not shown), confirming *a posteriori* the farm classification. All farms were positive for *Porcine Circovirus 2.* None of the animals investigated during the study displayed any clinical sign of these diseases.

Individual and collective medications administered to piglets were recorded during the study. None of the pigs received anti-microbial (AM) treatment until d26. During the first week after weaning, eight farms did not use any collective AM treatment (AM−, Table 1). In others (AM+), antimicrobials were administered via drinking water (1 farm), intramuscular injection (1 farm) or an AM supplemented starter diet (6 farms, Table 1).

### Animal data and sample collection

Piglets individual body weight (BW) was recorded on d26, d35 and d48. Faeces samples were collected on a sterile tube on d26 and d35, kept on ice until arrival at the laboratory, and then stored at −80°C until analysis. For practical reasons, faeces sampling were missing for one farm at d26 and one at d35. The occurrence of diarrhoeas in the batch of piglets was checked and registered daily by farmers.

The relative average daily gain (rADG) for the first 3 weeks after weaning was calculated for each pig: rADG = (weight at d48 – weight at d26) / N * 1 / weight at d26, N being the exact number of days between these two visits. Piglets were classified according to their rADG. In each farm, piglets showing the greater rADG (top 40% of the 18 sampled piglets distribution) and the lower rADG (bottom 40% of the distribution) were assigned to the rADG+ and rADG− classes respectively (approximately 7 piglets per class and per farm).

### Bacterial DNA extraction and sequencing

A modified version of the protocol by Godon *et al.* (1997) was used for DNA extraction. Briefly, after a lysis step by incubation for one hour at 70°C with Guanidine Thiocyanate and N-lauryl sarcosine, bacteria DNA was extracted using the chemagic STAR DNA BTS Kit for the automated isolation of DNA (Perkin Elmer, Wellesley, MA, USA). Quality and purity of the isolated DNA were checked by spectrophotometry on the NanoDrop (Fisher Scientific, Schwerte, Germany). The V3-V4 hypervariable regions of the 16S rRNA gene were amplified with the primers PCR1F_460: CTT TCC CTA CAC GAC GCT CTT CCG ATC TAC GGR AGG CAG CAG and PCR1R_460: GGA GTT CAG ACG TGT GCT CTT CCG ATC TTA CCA GGG TAT CTA ATCCT. The resulting PCR products were purified and loaded onto the Illumina MiSeq cartridge (Illumina Inc., San Diego, Ca, USA).

### Bioinformatics and biostatistics

The generated sequences were analysed using a subsampled open-reference OTU strategy with default settings in QIIME (v1.9.1)(Caporaso *et al.*, 2010). The paired-end reads were merged and demultiplexed. Subsampled open-reference OTU-picking was carried out using UCLUST with 97% sequence similarity. The taxonomy of each representative sequence was assigned using the UCLUST method on the Greengenes database with a 90% confidence threshold. Data chimera were checked using the Blast fragments approach (Altschul *et al.*, 1990) in QIIME. Low quality samples with less than 5 000 reads were excluded, leaving 222 samples from d26 and 254 samples from d35d, including 192 piglets having samples for both time points. Only OTUs of known phylum that were present in at least 5 samples, and had a relative abundance greater than 0.01% of total reads of the 476 samples were considered. These samples (minimum 4 831 reads) contained information relative to 1 177 OTUs (Supplementary Table 1).

The OTU table was imported in R computational language (R core team, ver. 3.3.2) for the ecological parameters evaluation. The variability within bacterial community (alpha diversity) and the differences among bacterial community (beta diversity) were calculated on the dataset rarefied to 4 831 reads per sample using the “phyloseq” (McMurdie and Holmes, 2013) and “Vegan” (Oksanen *et al.*, 2019) packages. Both qualitative (observed richness) and quantitative (Shannon index) alpha diversity indexes were taken into account. The influence of the investigated factors (age, farm, antimicrobial use, health status of the farm, growth class) on body weight and alpha diversity indexes were analysed by type 3 ANOVAs using the “lme4” (Bates *et al.*, 2015) and “car” (Fox and Weisberg, 2011) packages on R software. A first model including the farm (16 farms) and the interaction with age (d26 and d35) as fixed factors and the piglet as random factor was carried out. Then, a second model investigated simultaneously the effects of AM use (AM− vs. AM+) between weaning and d35, the health status of the farm (H+ vs. H−) and piglets growth class (rADG− vs. rADG+) as simple effects in interaction with age was performed. ADG and relative ADG were analysed in the same way, but without the age factor.

Regarding the beta diversity, the Unifrac distance matrix was calculated. The effect of farms, age, antimicrobial use and health status of the farm on the homogeneity of dispersion was determined using the betadisper function of Vegan. Then, a permutation analysis of variance (Adonis procedure with 999 permutations) including time, antimicrobial use and health status of the farm as factors was carried out. The farm factor was excluded by the permutational analysis since the betadisper result was significant, meaning that the null hypothesis (farms have the same dispersions) can be rejected (Anderson, 2006). The distance matrix was visualized with non-metric multidimensional scaling (NMDS) plots.

Relative abundances at family and genus level were calculated on unrarefied data. The methodology of Arumugam *et al.* (2011) was used to cluster piglets into groups of similar abundance called enterotypes. The clustering method was applied on the relative genus abundance. It used the Jensen-Shannon divergence distance and the Partitioning Around Medoids clustering algorithm. The optimal number of clusters was assessed using the Calinski-Harabasz Index. To evaluate the relevance of our enterotypes, the silhouette coefficient of our optimal clustering was compared to the silhouette coefficient of 100 of randomized dataset based on our dataset.

The influence of the investigated factors (age, farm, antimicrobial use, health status of the farm, growth class) on relative abundances was investigated using non parametric statistics. For the abundance of OTUs, the effects of age (d26 vs. d35), AM (AM− vs. AM+), H (H− vs. H+) and rADG (rADG+ vs. rADG−), were tested on the logarithmic fold-change ratios of taxa using a Benjamini-Hochberg (BH) correction to control the false discovery rate, with the “edgeR” package of R (Robinson *et al.*, 2010). For relative abundance aggregated at genus and family levels, the effects of factors with more than two levels (16 for the farm and 4 for the enterotype factor), were compared using Kruskal-Wallis and Dunn tests for each time point separately. The effect of two-level factors was first tested separately at d26 and d35, taking into account all individuals present for each time point, using, a Wilcoxon test with a BH correction. When the P-value was below 0.1 either at d26 or at d35, a model including the interaction of this factor with age was tested using the permutation tests of the lmPerm package of R software (Wheeler and Torchiano, 2016). For this, the dataset was restrained to individuals with data available for both time points. The effect of a factor was considered as significant if P < 0.05.

### Prediction of rADG using microbiota

The best linear model calculating rADG between d26 and d48 from microbiota data was determined using the Leaps package on R software (Lumley and Miller, 2017). The procedure was applied to a dataset including relative abundances of all families and genera, and of a preselection of OTUs identified as predictors of the rADG class at d26 and d35 as variables, and all piglets having data for both time-points (from 14 farms). These OTUs were identified using sparse Partial Least Square Discriminant Analysis with the MixOmics package (Le Cao *et al.*, 2016). To evaluate the robustness of the prediction, for each farm, coefficients were calculated from the dataset of the 13 other farms, were used to predict the rADG of piglets from the farm, and the R^2^ was calculated. The R^2^ presented in the result section is the mean adjusted-R^2^ of the 14 simulations corresponding to the 14 farms.

## Declarations

### Ethics approval

A competent ethics committee in animal experimentation has approved the experiment (authorization #CERVO-2016-6-V, committee of the national veterinary school of Nantes, France).

### Consent for publication

All authors have read and approved the final manuscript. The authors have no conflict of interests to declare.

### Availability of data and material

The raw reads obtained have been submitted to the Sequence Read Archive (SRA) of NCBI, under the BioProject ID PRJNA590802.

## Acknowledgements

To F. Thomas, M. Lefebvre, G. Martin, J.-Y. Audiard for their technical help and all pig producers who participated to this experiment. M. Leblanc-Maridor, N. Mach, Rogel-Gaillard, and P. Trevisi for their methodological and scientific advises.

## Supporting information

Additional Supporting Information may be found in a separate file on this web-site:

**Supplementary Table 1.** Description of the dataset.

**Supplementary Table 2.** Influence of age on the number of differentially expressed OTUs per family and on the relative abundance of each family.

**Supplementary Figure 1.** Distribution and clusters of samples from suckling 26-day old (n=222, pink dots) and weaned 35-day old piglets (n=254, blue dots) according to the non-metric multi-dimensional scaling (NMDS) on Unifrac distances calculated at the OTUs level.

**Supplementary Figure 2.** Proportions of OTUs in each family for which the abundance at d35 was significantly decreased (LogFC < 0, P < 0.05, n=514) or increased (LogFC > 0, P < 0.05, n=436) compared to d26. Faecal samples were collected from suckling 26-day old (n=222) and weaned 35-day old piglets (n=254) from 16 commercial farms. The numbers in brackets indicate the total number of OTUs present in each family.

**Supplementary Figure 3.** Farm variability in the response to weaning of the abundance of first 8 most abundant families of bacteria in faecal piglets samples. Samples were collected from suckling 26-day old (n=222) and weaned 35-day old piglets (n=254) from 16 commercial farms. Relative abundance of families from Firmicutes (A), as well as Bacteriodetes and Spirochaetes phyla (B) are presented. Among these families, before weaning, farm belonging influenced only the abundance of Christensenellaceae (P < 0.05). After weaning, the relative abundance of Ruminococcaceae, Lachnospiraceae, Clostridiaceae, Entrobacteriaceae and Spirochaetaceae were also different among farms (P < 0.05).

**Supplementary Figure 4.** Proportions of OTUs in each family for which the abundancy was significantly decreased (LogFC < 0, P < 0.05) or increased (LogFC > 0, P < 0.05) in low health status farms (H−) compared to farms with a satisfactory health status (H+). Faecal samples were collected from suckling 26-day old (A, n=222) and weaned 35-day old piglets (B, n=254) from 8 good health (H+) and 8 bad health (H−) commercial farms. The numbers in brackets indicate the total number of OTUs present in each family.

**Supplementary Figure 5.** Proportions of OTUs, in each family, for which the abundancy in AM+ animals was significantly decreased (LogFC < 0, P < 0.05, n=248) or increased (LogFC > 0, P < 0.05, n=214) compared to AM- animals. Faecal samples were collected from weaned 33-day old piglets from 16 commercial farms that received (AM+, n= 121) or not (AM-, n=133) a collective antimicrobial treatment between weaning (d27) and the day of sample collection. The numbers in brackets indicate the total number of OTUs present in each family.

**Supplementary Figure 6.** Proportions of OTUs, in each family, for which the abundancy in rADG- animals was significantly decreased (LogFC < 0, P < 0.05, n=150) or increased (LogFC > 0, P < 0.05, n=165) compared to rADG+ animals. Faecal samples were collected from suckling 26-day old (A) and weaned 33-day old piglets (B) from 16 commercial farms. In each farm, the 40% pigs showing the highest and 40% pigs showing the lowest relative ADG between d26 and d48 were attributed to high (rADG+, n=78 pigs with microbiota data at d26 and 100 at d35) or low (rADG-, n=88 pigs with microbiota data at d26 and 94 at d35) growth classes. The numbers in brackets indicate the total number of OTUs present in each family.

