## Supplementary for "The fecal microbiota of piglets is influenced by the farming environment and is associated with piglet robustness at weaning"

Additional Supporting Information for the manuscript:

1 Agricultural, Environmental, Food Science and Technology, Department of Agricultural and Food Sciences (DISTAL), Alma Mater Studiorum, University of Bologna, Italy.
2 PEGASE, INRAE, AGROCAMPUS OUEST, Saint Gilles, France.
3 Cooperl Arc Atlantique, Lamballe, France.
4 @BRIDGE, INRAE, Jouy-en-Josas, France.
5 GenPhySE, INRAE, Toulouse, France.
6 BIOEPAR, INRAE, ONIRIS, Nantes, France.

**Supplementary Table 1.** Description of the dataset.

| | Description | Nb of samples | Nb of OTUs | Minimal Nb of reads per sample |
|---|---|---|---|---|
| **26-day-old piglets** | Feces samples collected and extracted (including water and repeated samples) | 305 | 1504 | 0 |
| | Dataset including samples with at least 5 000 reads | 252 | 1504 | 5016 |
| | Dataset filtered by repeated samples and water samples | 222 | 1504 | 5016 |
| | Dataset including OTUs of known Phylum that are present in at least 5 samples | 222 | 1490 | 5003 |
| | Dataset after rarefaction (by resampling the OTU table such that all samples have the same library size) | 222 | 1490 | 5003 |
| | Dataset not rarefied filtered to keep OTUs with total abundance higher than 0.01% of the total number of reads | 222 | 1177 | 4831 |
| **35-day-old piglets** | Feces samples collected and extracted (including water and repeated samples) | 348 | 1504 | 1 |
| | Dataset including samples with at least 5 000 reads | 288 | 1504 | 5030 |
| | Dataset filtered by repeated samples and water samples | 254 | 1504 | 5030 |
| | Dataset including OTUs of known Phylum that are present in at least 5 samples | 254 | 1490 | 5030 |
| | Dataset after rarefaction (by resampling the OTU table such that all samples have the same library size) | 254 | 1490 | 5030 |
| | Dataset not rarefied filtered to keep OTUs with total abundance higher than 0.01% of the total number of reads | 254 | 1177 | 4958 |

Datasets after rarefaction are used to study alpha diversity.
Datasets not rarefied but filtered are used to do the other analyses.

**Supplementary Table 2.** Influence of age on the number of differentially expressed OTUs per family and on the relative abundance of each family.

OTUs were considered as influenced by AM treatmentage according to the level of the FDR of their logFold Change ratio.
Relative abundancies were calculated on row prefiltered data (no rarefaction nor TSS and CLR transformations).
The comparison between the two ages was performed by a Wilcoxon test with BH correction for each phylum / family.
The comparison between the 16 farms was performed by a Kruskal-Wallis test for each family.

| Abundance | | Nb OTUs | | | relative abundancy of the family (%) | | | P-value | | |
|---|---|---|---|---|---|---|---|---|---|---|
| rank | Family | in dataset | ↘ at d35 | ↗ at d35 | d26 + d35 | d26 | d35 | Time | farm at d26 | farm at d35 |
| 1 | f__Ruminococcaceae | 368 | 197 | 94 | 32.32 | 32.94 | 31.01 | 0.1485 | 0.331 | 0.000 |
| 2 | f__ | 201 | 93 | 47 | 15.31 | 17.39 | 13.98 | 0.002 | 0.053 | 0.001 |
| 3 | f__Prevotellaceae | 126 | 2 | 117 | 12.69 | 7.84 | 19.07 | <0.001 | 0.624 | 0.100 |
| 4 | f__Lachnospiraceae | 178 | 60 | 91 | 12.25 | 10.82 | 13.13 | <0.001 | 0.221 | 0.026 |
| 5 | f__Christensenellaceae | 35 | 31 | 1 | 4.51 | 5.55 | 3.63 | <0.001 | 0.036 | 0.018 |
| 6 | f__Bacteroidaceae | 38 | 30 | 2 | 3.95 | 5.33 | 2.07 | <0.001 | 0.205 | 0.056 |
| 7 | f__S24-7 | 34 | 11 | 16 | 2.56 | 2.08 | 2.46 | <0.001 | 0.205 | 0.001 |
| 8 | f__Enterobacteriaceae | 4 | 3 | | 2.41 | 2.99 | 1.74 | <0.001 | 0.052 | 0.000 |
| 9 | f__Clostridiaceae | 21 | 10 | 8 | 1.92 | 1.97 | 1.47 | <0.001 | 0.055 | 0.000 |
| 10 | f__Spirochaetaceae | 27 | 7 | 12 | 1.74 | 1.86 | 1.76 | 0.0712 | 0.302 | 0.000 |
| 11 | f__[Paraprevotellaceae] | 27 | 3 | 20 | 1.43 | 1.09 | 2.16 | <0.001 | 0.536 | 0.000 |
| 12 | f__Porphyromonadaceae | 20 | 12 | 1 | 1.40 | 1.81 | 0.87 | <0.001 | 0.140 | 0.000 |
| 13 | f__Fusobacteriaceae | 6 | 6 | | 1.34 | 2.05 | 0.79 | <0.001 | 0.501 | 0.218 |
| 14 | f__Lactobacillaceae | 10 | 7 | 1 | 0.96 | 0.98 | 0.75 | 0.0119 | 0.032 | 0.001 |
| 15 | f__Campylobacteraceae | 7 | 2 | 4 | 0.94 | 0.81 | 1.11 | 0.1007 | 0.042 | 0.040 |
| 16 | f__Succinivibrionaceae | 10 | 1 | 7 | 0.77 | 0.60 | 1.12 | 0.0071 | 0.777 | 0.000 |
| 17 | f__p-2534-18B5 | 1 | 1 | | 0.64 | 0.90 | 0.56 | <0.001 | 0.655 | 0.000 |
| 18 | f__Erysipelotrichaceae | 9 | 5 | 1 | 0.50 | 0.45 | 0.36 | 0.4766 | 0.159 | 0.256 |

| | | | | | | | | | | |
|---|---|---|---|---|---|---|---|---|---|---|
| 19 | f__Desulfovibrionaceae | 6 | 6 | | 0.41 | 0.48 | 0.27 | <0.001 | 0.256 | 0.292 |
| 20 | f__RF16 | 5 | 1 | 1 | 0.35 | 0.29 | 0.39 | 0.1485 | 0.092 | 0.010 |
| 21 | f__[Odoribacteraceae] | 7 | 7 | | 0.30 | 0.42 | 0.14 | <0.001 | 0.299 | 0.460 |
| 22 | f__Veillonellaceae | 7 | | 7 | 0.23 | 0.11 | 0.31 | <0.001 | 0.488 | 0.027 |
| 23 | f__Alcaligenaceae | 6 | 3 | 1 | 0.19 | 0.18 | 0.15 | 0.0648 | 0.401 | 0.179 |
| 24 | f__Helicobacteraceae | 2 | 1 | | 0.18 | 0.17 | 0.16 | 0.0944 | 0.386 | 0.000 |
| 25 | f__[Mogibacteriaceae] | 5 | 4 | | 0.14 | 0.16 | 0.10 | <0.001 | 0.012 | 0.135 |
| 26 | f__Peptostreptococcaceae | 2 | | 2 | 0.11 | 0.07 | 0.14 | <0.001 | 0.063 | 0.007 |
| 27 | f__Enterococcaceae | 2 | 2 | | 0.11 | 0.12 | 0.01 | <0.001 | 0.363 | 0.166 |
| 28 | f__Streptococcaceae | 2 | 1 | 1 | 0.10 | 0.17 | 0.07 | 0.1114 | 0.001 | 0.000 |
| 29 | f__BS11 | 1 | 1 | | 0.071 | 0.075 | 0.060 | 0.4186 | 0.271 | 0.166 |
| 30 | f__Peptococcaceae | 2 | 1 | | 0.053 | 0.054 | 0.027 | 0.0365 | 0.020 | 0.025 |
| 31 | f__Rikenellaceae | 2 | 2 | | 0.046 | 0.085 | 0.016 | 0.0317 | 0.421 | 0.262 |
| 32 | f__Pasteurellaceae | 4 | 2 | 2 | 0.028 | 0.055 | 0.045 | <0.001 | 0.639 | 0.163 |
| 33 | f__Turicibacteraceae | 1 | 1 | | 0.019 | 0.014 | 0.010 | <0.001 | 0.262 | 0.006 |
| 34 | f__Oxalobacteraceae | 1 | 1 | | 0.012 | 0.005 | 0.006 | 0.0140 | 0.522 | 0.019 |
| | Total | 1177 | 514 | 436 | 100.00 | 100 | 100 | | | |

**Supplementary Figure 1.** Distribution and clusters of samples from suckling 26-day old (n=222, pink dots) and weaned 35-day old piglets (n=254, blue dots) according to the non-metric multi-dimensional scaling (NMDS) on Unifrac distances calculated at the OTUs level.

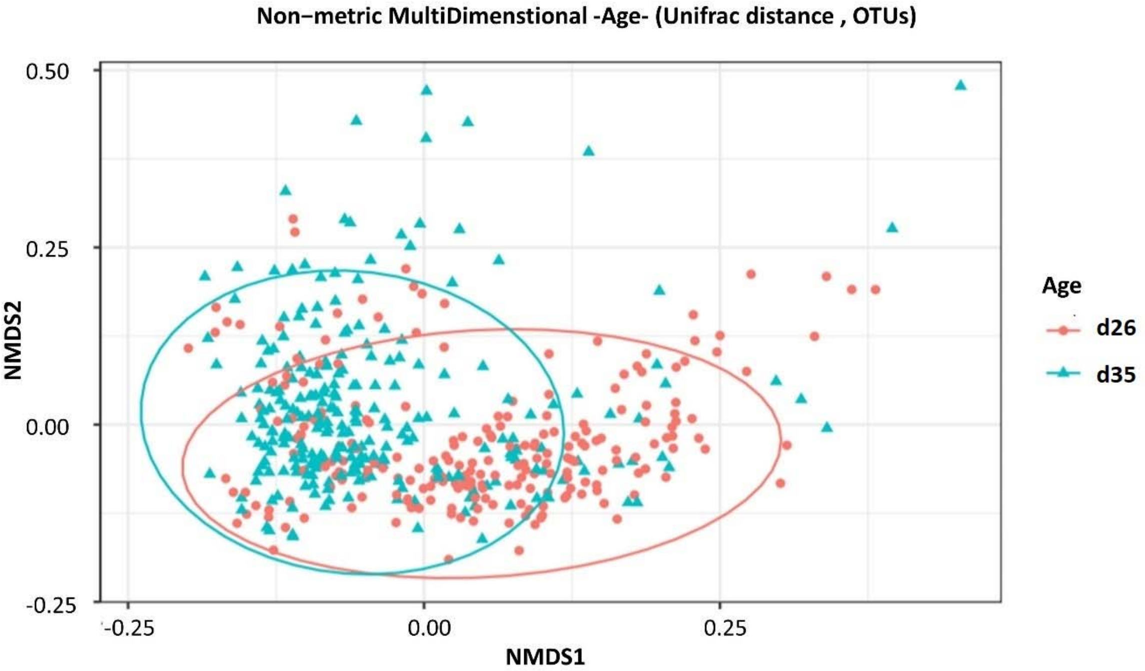


**Supplementary Figure 2.** Proportions of OTUs in each family for which the abundance at d35 was significantly decreased (LogFC < 0, P < 0.05, n=514) or increased (LogFC > 0, P < 0.05, n=436) compared to d26. Faecal samples were collected from suckling 26-day old (n=222) and weaned 35-day old piglets (n=254) from 16 commercial farms. The numbers in brackets indicate the total number of OTUs present in each family.

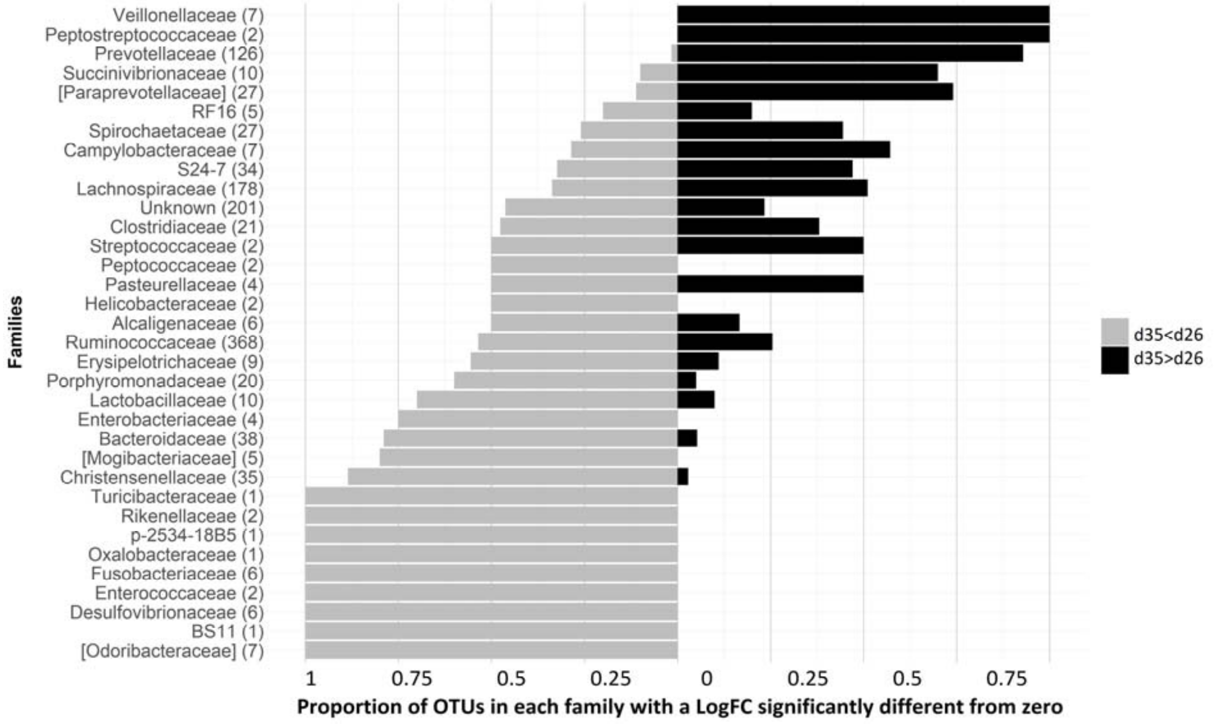

**Supplementary Figure 3.** Farm variability in the response to weaning of the abundance of first 8 most abundant families of bacteria in faecal piglets samples. Samples were collected from suckling 26-day old (n=222) and weaned 35-day old piglets (n=254) from 16 commercial farms. Relative abundance of families from Firmicutes (A), as well as Bacteriodetes and Spirochaetes phyla (B) are presented. Among these families, before weaning, farm belonging influenced only the abundance of Christensenellaceae (P < 0.05). After weaning, the relative abundance of Ruminococcaceae, Lachnospiraceae, Clostridiaceae, Entrobacteriaceae and Spirochaetaceae were also different among farms (P < 0.05).

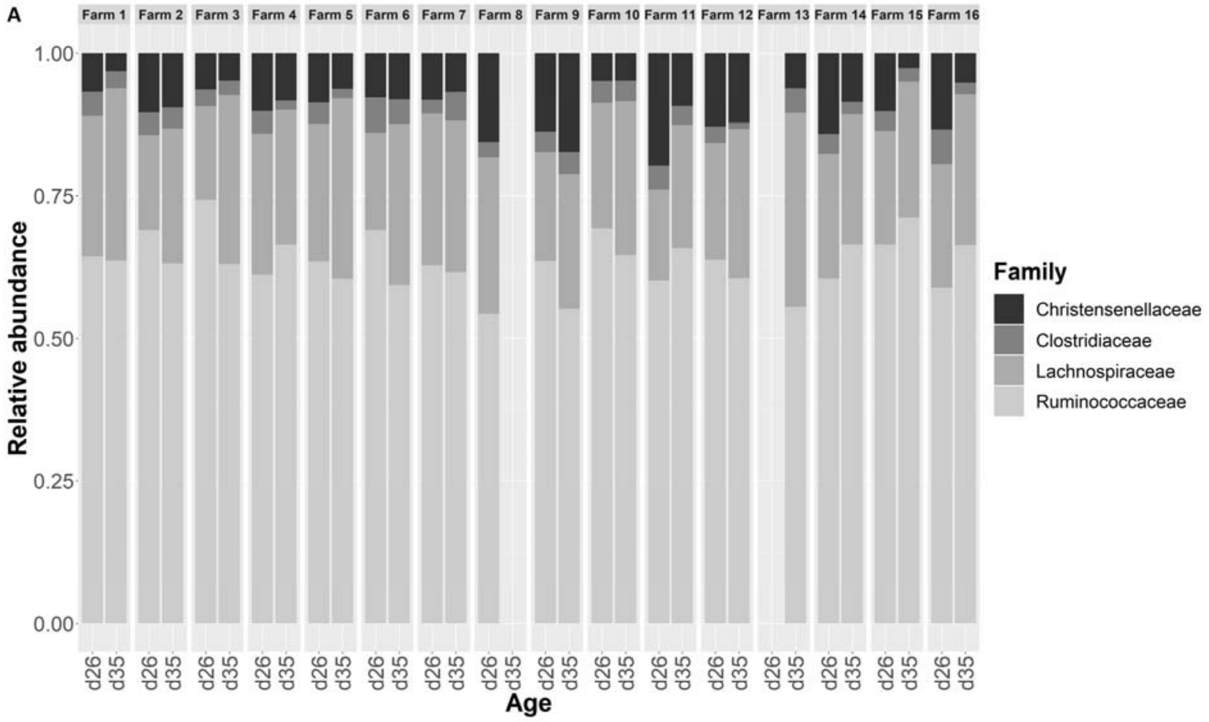


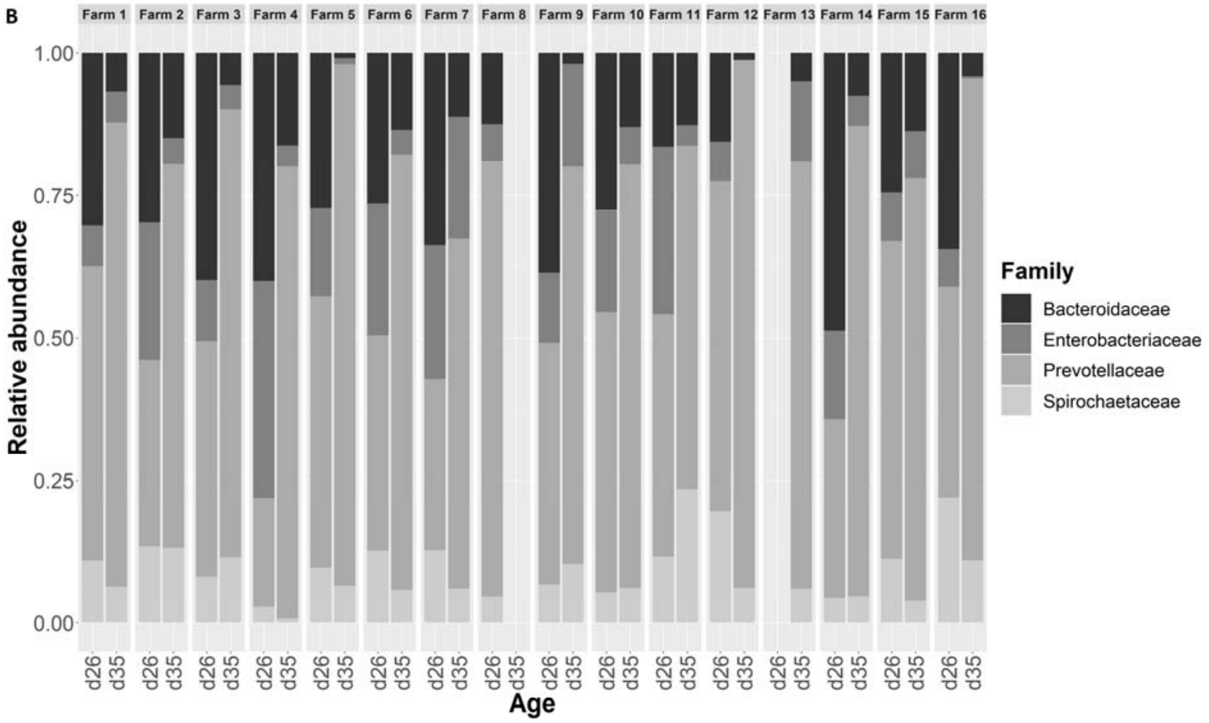

**Supplementary Figure 4.** Proportions of OTUs in each family for which the abundancy was significantly decreased (LogFC < 0, P < 0.05) or increased (LogFC > 0, P < 0.05) in low health status farms (H-) compared to farms with a satisfactory health status (H+). Faecal samples were collected from suckling 26-day old (A, n=222) and weaned 35-day old piglets (B, n=254) from 8 good health (H+) and 8 bad health (H-) commercial farms. The numbers in brackets indicate the total number of OTUs present in each family.

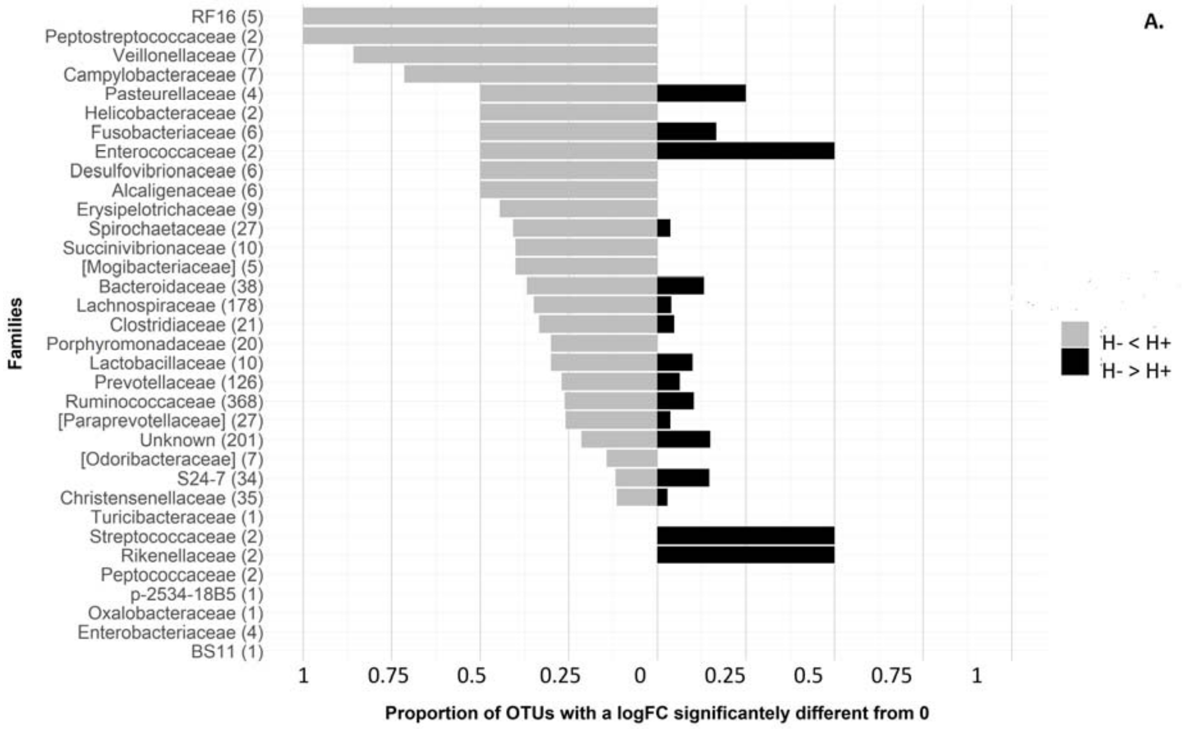


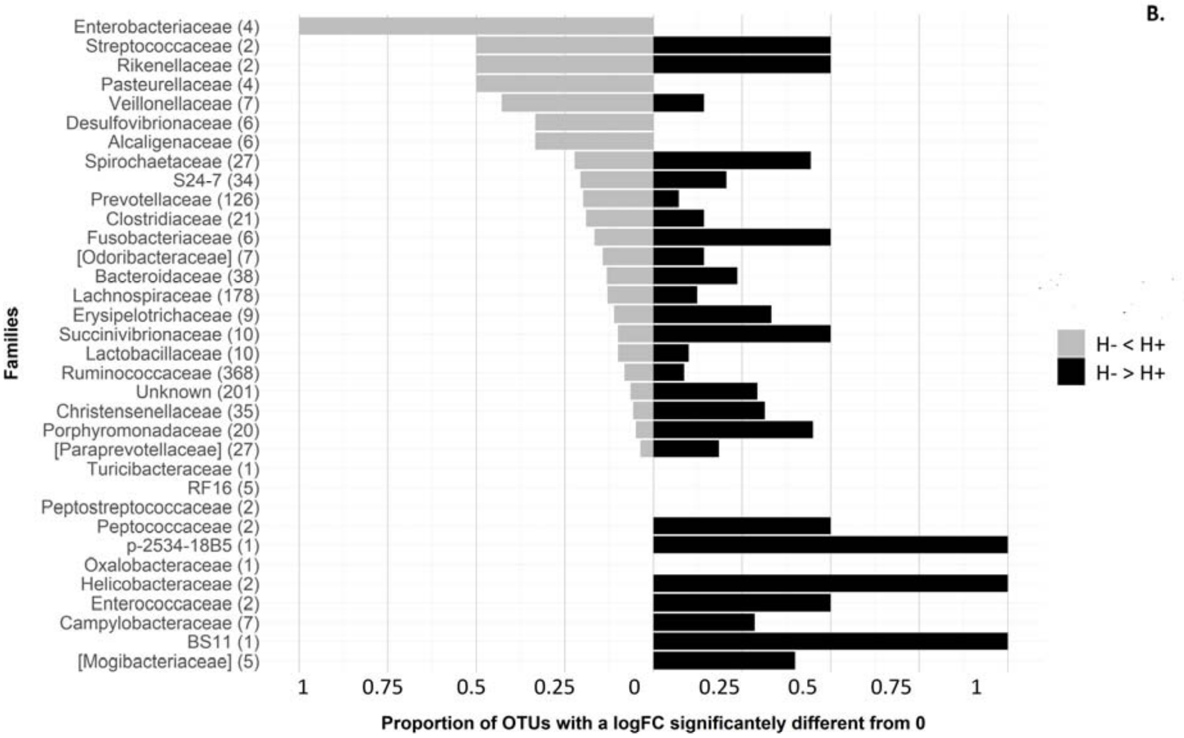

**Supplementary Figure 5.** Proportions of OTUs, in each family, for which the abundancy in AM+ animals was significantly decreased (LogFC < 0, P < 0.05, n=248) or increased (LogFC > 0, P < 0.05, n=214) compared to AM- animals. Faecal samples were collected from weaned 33-day old piglets from 16 commercial farms that received (AM+, n= 121) or not (AM-, n=133) a collective antimicrobial treatment between weaning (d27) and the day of sample collection. The numbers in brackets indicate the total number of OTUs present in each family.

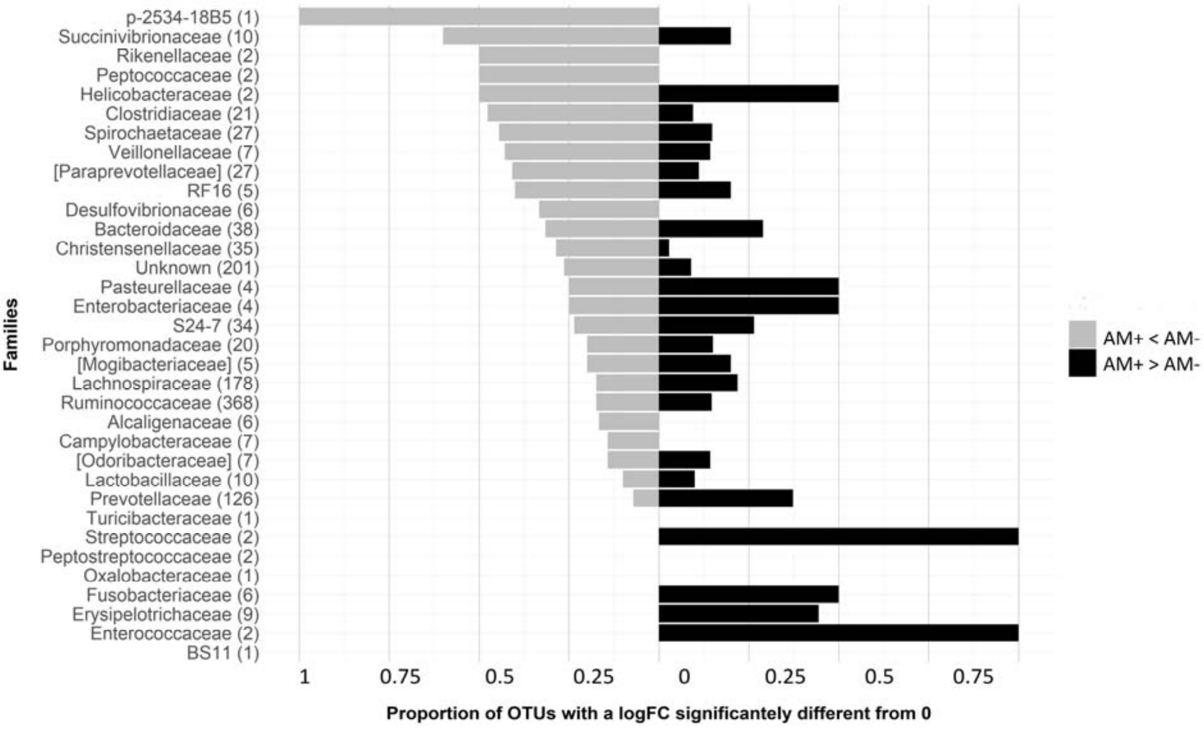

**Supplementary Figure 6.** Proportions of OTUs, in each family, for which the abundancy in rADG- animals was significantly decreased (LogFC < 0, P < 0.05, n=150) or increased (LogFC > 0, P < 0.05, n=165) compared to rADG+ animals. Faecal samples were collected from suckling 26-day old (A) and weaned 33-day old piglets (B) from 16 commercial farms. In each farm, the 40% pigs showing the highest and 40% pigs showing the lowest relative ADG between d26 and d48 were attributed to high (rADG+, n=78 pigs with microbiota data at d26 and 100 at d35) or low (rADG-, n=88 pigs with microbiota data at d26 and 94 at d35) growth classes. The numbers in brackets indicate the total number of OTUs present in each family.

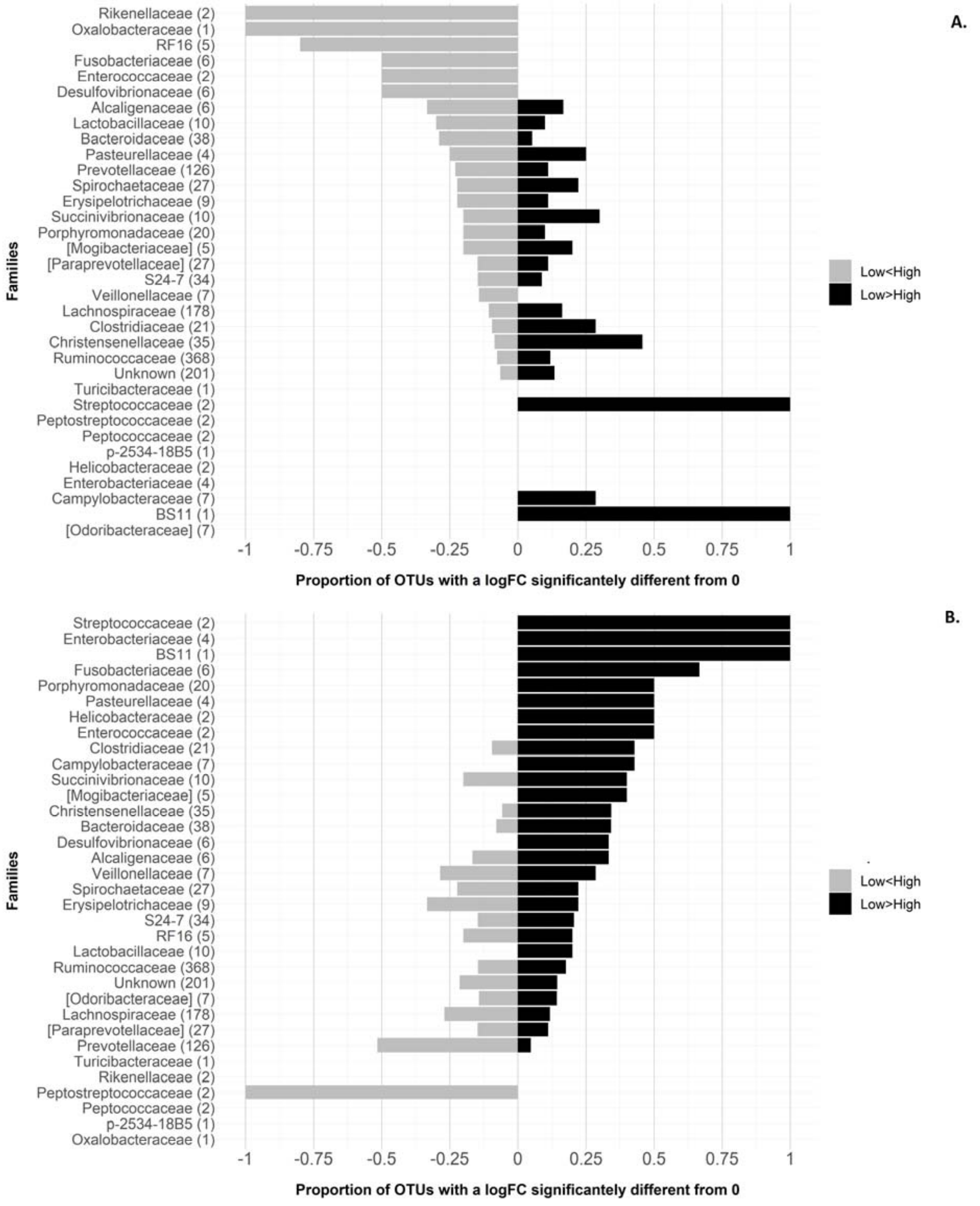
